# Malthusian Catastrophe: Species Extinction Caused by Oversized Population

**DOI:** 10.1101/134221

**Authors:** Xubin Pan

**Affiliations:** Institute of Plant Quarantine, Chinese Academy of Inspection and Quarantine, Beijing 100029, China

**Keywords:** Allee threshold, Pan threshold, carrying capacity, disturbance, species immigration

## Abstract

There is one pseudo-extinction debt and four occurring conditions for real extinction debt. Since small and oversized populations have a high extinction risk, Pan threshold (upper limit) was calculated for Verhulst-Pear “logistic” growth model and logistic model with the Allee effect, an important parameter corresponding to Allee threshold (lower limit).

---

The prediction and management of population dynamic is central to human sustainable development and ecosystem protection. We have been aware that small population has a high risk of extinction rate [1] and our society has utilized many resources to save endangered species. However, if we overlooked the extinction risk caused by oversized population, Malthusian catastrophe would happen [2].

The extinction of most species was a long-term process, and few species suddenly became extinct. The phenomenon has been coined as extinction debt in ecology, which is worth investigating due to its application in wildlife conservation and invasive species management. Particularly, species extinction rate has been accelerated due to climate change [3]. If we understand well the occurring conditions of extinction debt, we will be able to slow down the extinction of or even save endangered species, or eliminate undesired alien species in someplace.

Several studies are available on the mechanism of extinction debt caused by changes in habitat quality/quantity/connectivity [4,5]. Habitat loss or degradation might repel one species and favor another. Furthermore, little research has been conducted on the link between habitat change and population change embedded in the context of extinction debt. Thus, it is necessary to explore the occurring conditions of extinction debt for specific species and how the extinction debt will be affected by forcing events. Unfortunately, few records have been found on the occurring conditions taking into account population dynamics. To simplify population dynamics, we proposed a framework with the fluctuation of population size (*N*) around carrying capacity (*K*) with self-adjustment as a quasi-steady state. If the population size falls below the Allee threshold (*A*), extinction debt will occur, the process of which will be irreversible. Normally, the species’ population can sustain continuously until the population and/or the habitat are highly impacted by external disturbances.

One type of direct extinction occurs when a species disappear immediately due to the disturbance at time *t*, when the population size is decreased to 0 (Figure 1A, *N_t_*=0). As for another type of direct extinction or pseudo-extinction debt, the species population can sustain for a short period of time after the disturbance, since population does not have any new individuals but old, injured or gender-imbalanced individuals. In this case, the time interval from *N_t_* to 0 (extinction) is shorter than the life span of the species (Figure 1B, *N_t_*>0).

**Figure 1.**
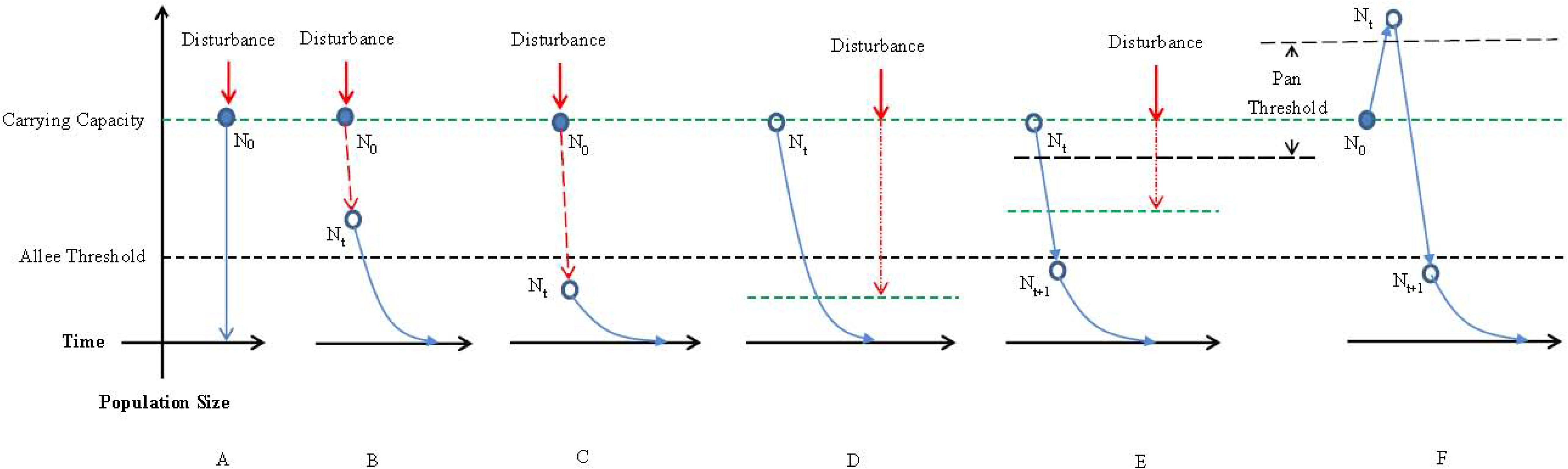
The occurring conditions of extinction debt in a habitat due to the disturbance impact in time *t* on the population size *N0* (Fig. 1A, 1B and 1C), carrying capacity (Fig. 1D and 1E) and population dynamics (Fig. 1F).

For the occurrence of extinction debt, there are two typical conditions. One is that *N_0_* decreases to *N_t_* due to the disturbance at time *t*, which is less than *A* (Figure 1C). The other is that *K* shrink to less than *A* due to the disturbance at time *t* (Figure 1D). Then *N_t_* follows *K* and falls below *A* eventually.

When the population size is larger than *K*, it will decrease or regress to *K*, leading to fluctuation of *N* around *K* under normal conditions. If the population size is too large (here we define the existing limit as the Pan threshold, *P*), the call-back of *N* can be zero or less than *A*, resulting in an extinction debt finally. Two conditions might lead to such an outcome. One condition is that *K* is suppressed by the disturbance so that *N_t_* is higher than *P* (Figure 1E, *P* is a function of *K* and positively correlated with *K*. The other is that *N_0_* is raised to *N_t_* by external force, which is higher than *P* (Figure 1F).

The formula for calculating Pan threshold (*P*) of the two typical population growth models, Verhulst-Pear “logistic” growth model [6] and logistic model with the Allee effect [7], is as follows:

For the Verhulst-Pear “logistic” growth model,

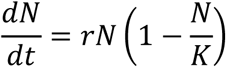

where *r* is the growth rate. When 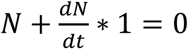, we get the Pan threshold

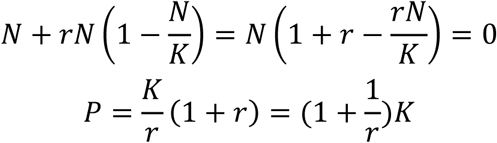

For the logistic model with the Allee effect,

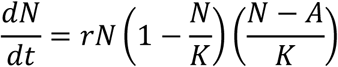

When 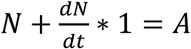, we get the Pan threshold

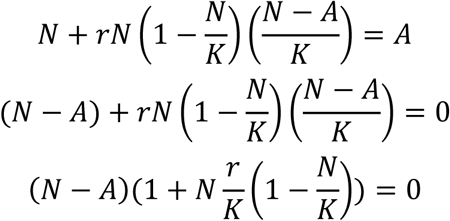

*N>A* and *K*>0, 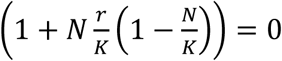

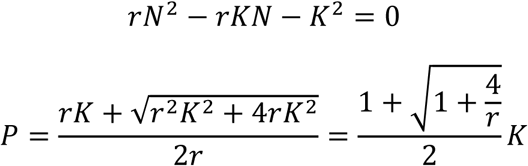

For both growth models, *P* increases with the increase of *K* or the decrease of *r*. And *P* is not relevant to *A* in both models.

There are two dead zones for species population, meaning either small population or oversized population can lead to species extinction. One is below the Allee threshold, and the other is above the Pan threshold. If the population size falls within these two dead zones, the species will be in the process of extinction debt. Thus, appropriate population size should be within [*A*, *P*] (Figure 2a). Both *A* and *K* are equilibrium points, and the point *P* will transform to *A* soon. Others points in the interval will fluctuate around the point *K*. Both extremely small (*N*<*A*) and extremely large (*N*>*P*) population will go extinct eventually. Thus we can adjust the population (*r*) or carrying capacity (*K*) to control the occurrence of extinction debt according to policy goals. Species immigration, which is not considered in the above simplified frame, also exerts a significant impact on the extinction debt. Based on our analysis, in Figure 1C, species immigration can help increase the population size and produce a credit, especially when considering the effect of genetic exchange of species on the Allee threshold (*A*) [4]. However, Figure 1E and 1F reveal that, species immigration can become a burden for existing large-sized population.

**Figure 2.**
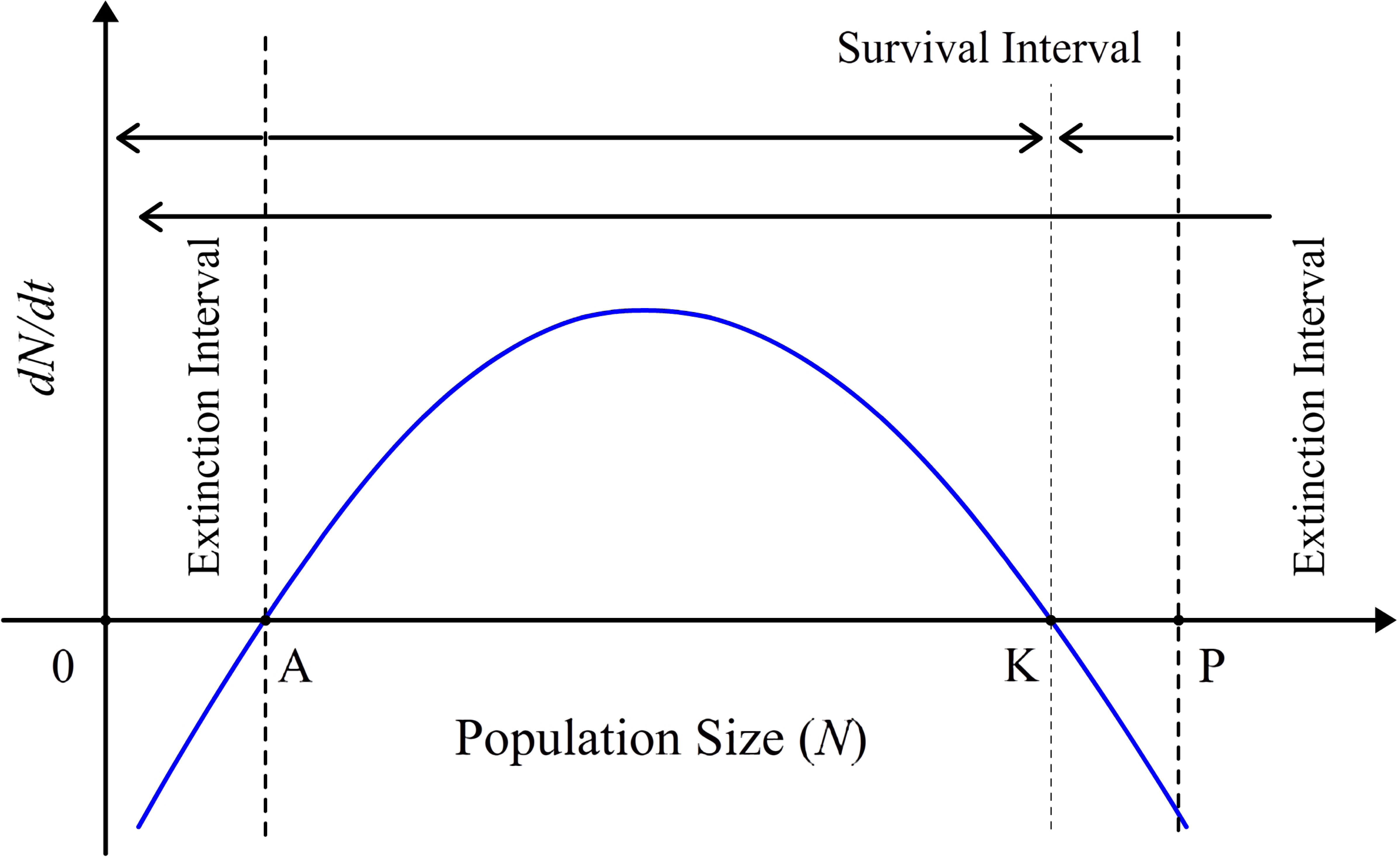
Allee threshold (*A*), carrying capacity (*K*) and Pan threshold (P) (2A). The population size can be divided into three sections: survival interval and two extinction intervals (2A). Population dynamics of prey and predator species (2B). There is a lag between the prey population peak and the predator population peak (2B).

Therefore, the Allee threshlod [8], or the Pan threshold can be utilized to improve endangered species conservation and invasive species control. For example, there is a lag between the peaks of the two species in the prey-predator (Lotka–Volterra) equations [9] (Figure 2b). Allee effect can be employed to control the predator population directly, while Pan threshold to regulate the prey population (*K*). When the predator population size is large, and the prey population is continuously decreased due to natural or anthropogenic disturbances, the population of predator can easily surpass the Pan threshold and become extinct. This might be the possible mechanism of extinction for many flourishing species in the geological history.

## Conclusions

There is one pseudo-extinction debt and four occurring conditions for real extinction debt in the frame with Allee threshold, carrying capacity and exogenous disturbance. For Verhulst-Pear “logistic” growth model and logistic model with the Allee effect, Pan threshold (upper limit) is proposed as an important parameter corresponding to Allee threshold (lower limit). The measures considering the occurring conditions of extinction debt would further improve biodiversity conservation and bio-invasion control.

## Declarations section

*Ethics approval and consent to participate*

Not applicable.

*Consent for publication*

All author agrees this submission. *Availability of data and material* Not applicable.

*Funding*

The work is supported by the Beijing NOVA Programme (Z1511000003150107).

### Acknowledgments

We warmly thank Fengqiao Liu for helpful suggestions for the manuscript revision.

## Conflict of Interest

The authors declare that they have no conflict of interest.

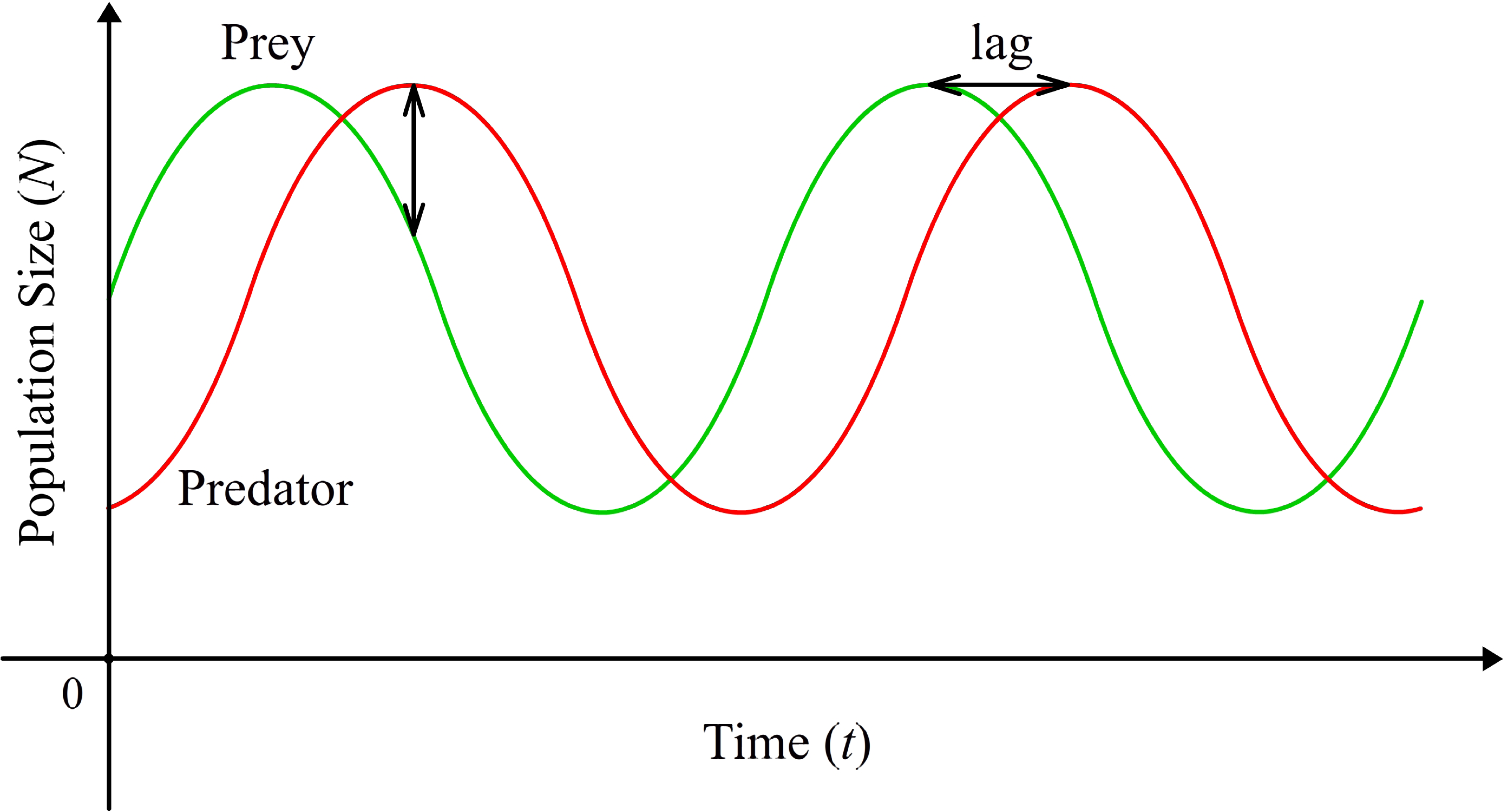

